# Ecological dynamics of the almond floral microbiome in relation to crop management and pollination

**DOI:** 10.1101/2020.11.05.367003

**Authors:** Robert N. Schaeffer, David W. Crowder, Javier Gutiérrez Illán, John J. Beck, Tadashi Fukami, Neal M. Williams, Rachel L. Vannette

**Affiliations:** Department of Entomology, Washington State University, Pullman, WA 99164; Department of Entomology & Nematology, University of California Davis, Davis, CA 95616; Chemistry Research Unit, Center for Medical, Agricultural, and Veterinary Entomology, Agricultural Research Service, United States Department of Agriculture, Gainesville, FL 32608; Department of Biology, Stanford University, Stanford, CA 94305

**Keywords:** agricultural landscapes, ecosystem services, flower microbiome, fungicide, pollination, *Prunus dulcis*, sustainable agriculture

## Abstract

1. Crop tissues harbor microbiomes that can affect host health and yield. However, processes driving microbiome assembly, and resulting effects on ecosystem services, remain poorly understood. This is particularly true of flowering crops that rely on pollinators for yield.
2. We assessed effects of orchard management tactics and landscape context on the flower microbiome in almond, *Prunus dulcis*. Fourteen orchards (5 conventional, 4 organic, 5 habitat augmentation) were sampled at two bloom stages to characterize bacterial and fungal communities associated with floral tissues. The surveys were complemented by *in vitro* experiments to assess effects of arrival order and fungicides on nectar microbial communities, and effects of fungicides and microbes on honey bee foraging. Finally, a field trial was conducted to test effects of fungicides and microbes on pollination.
3. As bloom progressed, bacterial and fungal abundance and diversity increased, across all floral tissue types and management strategies. The magnitude by which microbial abundance and diversity were affected varied, with host proximity to apiaries and orchard management having notable effects on bacteria and fungi, respectively.
4. Experiments showed immigration history and fungicides affected the composition of nectar microbial communities, but only fungicides affected pollinator foraging through reduced nectar removal. Neither treatment affected pollination services.
5. *Synthesis and applications*. Our results shed light on routes through which management practices can shape microbiota associated with flowers of a pollinator-dependent crop. With growing appreciation for the role of floral-associated microbes in affecting biotic interactions at the floral interface, understanding such drivers can potentially inform microbial-derived ecosystem services in agricultural landscapes, including pollination and biocontrol.

## Introduction

Crop tissues harbor distinct microbiomes that affect host health and yield. For example, microbiomes can affect host tolerance to stress and disease (Vurukonda *et al*., 2016; Berg and Koskella, 2018) and can stimulate growth through mobilization and transport of nutrients (Pii *et al*., 2015). However, despite the importance of the microbiome, few studies have investigated the processes mediating microbiome assembly, especially in crop systems (but see Edwards *et al*., 2015; Grady *et al*., 2019). Assessing such processes is critical, as manipulation of the plant microbiome can have wide-ranging effects, including improved crop performance, food safety, associated ecosystem services, and sustainability (Mueller and Sachs, 2015; Busby *et al*., 2017; Toju *et al*., 2018; Allard and Micallef, 2019).

Approximately 35% of crops produced globally benefit from pollination by arthropods (Klein *et al*., 2007). With an economic value exceeding $300 billion globally (Lautenbach *et al*., 2012), there is strong incentive to manage pollination of flowering crops during bloom. Bloom management is also critical because some plant pathogens infect crops through flowers and can be dispersed by pollinators (McArt *et al*., 2014). To combat pathogens, growers often use fungicides, bactericides, or other antibiotics during bloom. However, such chemicals can cause non-target effects on the microbiome (Schaeffer *et al*., 2017) and affect pollinators (Frazier *et al*., 2015; Johnson, 2015).

In addition, agricultural intensification often decreases native pollinator abundance and diversity, largely through habitat loss (Kremen *et al*., 2002; Kovács□Hostyánszki *et al*., 2017; Grab *et al*., 2019). Given that pollinators harbor distinct microbiomes and disperse microbes between flowers, loss of pollinator biodiversity and reduced pollinator visitation may indirectly affect microbiome assembly (Vannette and Fukami, 2017; Russell *et al*., 2019). In orchard systems, flowering strips, hedgerows, and nesting structures have been employed to support pollinator populations (Scheper *et al*., 2015; Williams *et al*., 2015; Kremen *et al*., 2019). However, whether restored vegetation, which can act as a source of inocula, affects the assembly of microbiomes in crop systems remains poorly understood (Lindow and Andersen, 1996; Lymperopoulou *et al*., 2016).

Here, we assessed how agricultural management affected flower microbiome assembly and function in mass-flowering almond (*Prunus dulcis*). Almond depends on biotic pollination for fruit set, and is heavily managed during bloom to ensure adequate pollination and prevention of pathogens, namely *Monilinia laxa*, the causal agent of brown rot blossom blight. Synthetic fungicides are applied during bloom to preempt *M. laxa* establishment in conventional orchards (Adaskaveg *et al*., 2017). However, increasing demand for sustainably produced almonds has spurred adoption of organic management tactics in many orchards (Brodt *et al*., 2009; Plattner *et al*, 2013), including use of copper and other materials instead of synthetics for disease control. Given that pollinators of almond, including honey bees and bumble bees, are sensitive to the chemical alterations that nectar microbes induce through metabolism of nectar resources (Rering *et al*., 2018; Schaeffer *et al*., 2019), shifts in microbiome structure arising from different management schemes may affect pollination services (Herrera *et al*., 2013; Vannette *et al*., 2013; Schaeffer and Irwin, 2014).

To address these linkages, we conducted a field survey of microbial diversity on almond flowers in orchards with different management schemes. We also conducted two experiments to examine mechanisms that affect microbial assembly and function in almond. First, we examined how fungicides interact with variation in microbial immigration history to affect microbial community structure. Second, we examined effects of nectar microbes and fungicides on honey bee foraging, and consequences for pollination. Overall, our results provide evidence that variation in management of a blooming crop can shape microbiome assembly and the biotic interactions that mediate ecosystem services in agricultural landscapes.

## Methods

### Study system

California (CA) almonds flower in February through early March, and rely almost exclusively on managed *Apis mellifera* colonies for pollination. Approximately 470,000 ha of almond orchards in CA produced over 80% of the world’s supply in 2018, generating $5.8B in farmer revenue (Sumner *et al*., 2014; Almond Board of California, 2019). Despite these numbers, fruit set in almond orchards typically ranges from 10-40% (Bosch and Blas, 1994), partly due to limited pollinator availability and the quality of services they provide, in addition to disease and inclement weather during bloom.

### Orchard survey

We surveyed 14 orchards (4 organic, 5 conventional, and 5 with supplemental forb plantings) across the Sacramento Valley of CA (Figure 1). Supplemental forb plantings included a mix of annual species native to CA, including *Calandrinia ciliata, Collinsia heterophylla, Eschscholzia californica, Nemophila maculata, Nemophila menziesii, Phacelia campanularia*, and *Phacelia ciliata*. Beyond the addition of forb plantings, forb-amended orchards follow conventional management practices. Orchards were sampled twice between February 15 and 24, 2017, once at early bloom (~10% of flowers open) and then at peak bloom (>50% of flowers open). At each orchard and sampling event, six trees (‘Nonpareil’ variety) were sampled; three near the edge of the orchard, and adjacent to the forb planting if available. These trees were located in the second row in from the edge of the orchard (‘edge’). The other three trees were sampled from the orchard interior (row 10) (‘interior’). We chose this sampling scheme because semi-natural habitat in the surrounding landscape can increase visitation by native pollinators such as bees and flies (Klein *et al*., 2012). Pollinators can also be important dispersal agents for microbes (Aizenberg-Gershtein *et al*., 2013; Vannette and Fukami, 2017); thus, we may see greater microbial abundance or diversity in flowers in close proximity to these natural habitats. For each site (edge or interior) and sampling event, 30 open flowers (*N* = 10 per tree) were collected using aseptic technique and then pooled at the site level. Flowers with flat, fully-reflexed petals that had been open for approximately 3 days were chosen for collection (Yi *et al*., 2006). This choice increased the probability that flowers had been visited by pollinators that disperse microbes. Once collected, flowers were placed in a cooler and transferred to the lab, then stored at 4 °C until processing (within the following 24 h).

**Figure 1.**
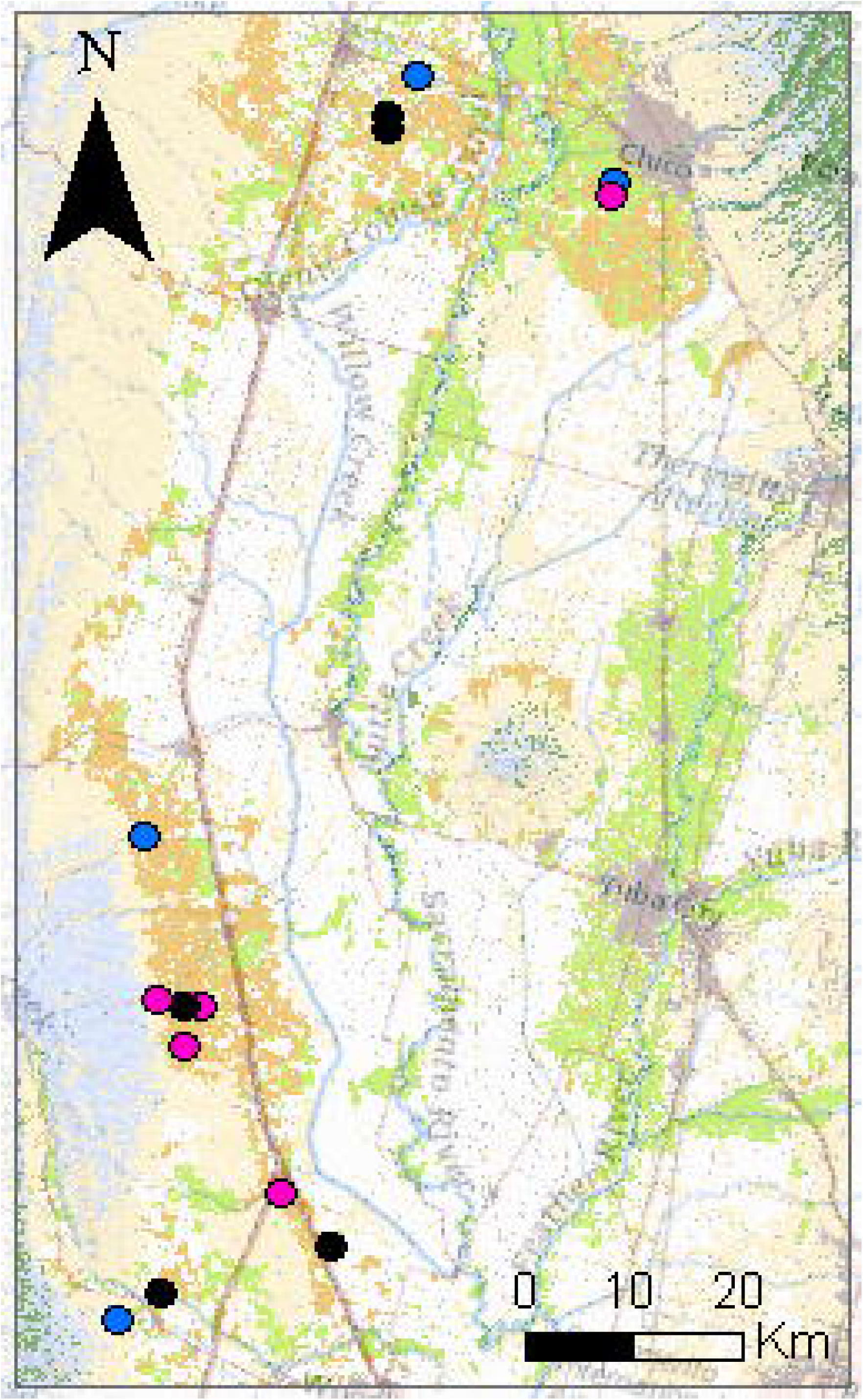
Map of the study area, showing heterogeneity in land-cover classes (Almond: brown, Forest: green; Shrubland: blue; Grass/Pasture: tan) across the Sacramento Valley of California, a major production area for almond. Points represent sampled orchards and are color-coded by management scheme (Conventional: black; Forb-amended: purple; Organic: blue).

To assess effects of pollinator foraging activity on microbiome structure and diversity, we used two complementary approaches. First, during flower collection, we measured the distance between trees sampled and the nearest set of honeybee hives in the orchard. Honey bees often forage near their hive (Gary *et al*., 1978), and we hypothesized that tree proximity would be a proxy for visitation frequency during bloom. Second, we measured the amount of semi-natural habitat within 1 km buffer of each orchard, as prior work shows increased semi-natural habitat can increase the diversity and visitation rates of native pollinators in orchards (Klein *et al*., 2012). We classified land cover from the croplands data layer product (USDA, 2017) within a 1-km radius of each orchard edge using ArcGIS (ESRI, Redlands, CA, USA). Natural habitat near orchards in this study can primarily be classified as chaparral, oak woodland, or valley and foothill riparian woodland (Barbour *et al*., 2007). We predicted that more natural habitat would promote more diverse floral microbiomes as pollinator species harbor distinct microbiomes (Koch *et al*., 2013; Graystock *et al*., 2017) and disperse microbes (Vannette and Fukami, 2017; Russell *et al*., 2019).

### Sample processing

Whole flowers were dissected acropetally in sequence to minimize cross-contamination, as previous work has shown that floral microbial communities can display taxonomic-structuring across tissue types (Junker and Keller, 2015). Petals were removed first using sterile forceps. Petals from all 30 flowers in a sample were then pooled in a 50 mL Falcon tube (Corning, Corning, NY, USA), massed for fresh weight (g), then suspended in 20 mL of 1x-0.15% PBS-Tween solution. The androecium and gynoecium (hereafter collectively referred to as ‘Anthers’) were then removed from the base of each flower, pooled in a 15 mL Falcon tube, massed, then suspended in 5 mL of 1x-0.15% PBS-Tween solution. To sample nectar, we ‘washed’ each hypanthia with 2 μL of 1x-0.15% PBS-Tween solution using a pipette, and pooled for each set of 30 flowers. Each wash was diluted with 1 mL of 1x-0.15% PBS-Tween solution. For petals, androecium, and gynoecium, samples were sonicated (Branson CPX5800H, Danbury CT) for 10 min to dislodge epiphytic microbes. After sonication, debris was removed from sample tubes by pouring samples through autoclaved cheesecloth into a sterile Falcon tube. Falcon tubes containing debris-filtered samples were then centrifuged at 3000 rpm for 10 min at 4 °C to pellet microbial cells. We then decanted the supernatant, re-suspended microbial cell pellets in 1 mL of autoclaved PBS solution, vortexed tubes, then transferred the cell suspensions to new 1.7 mL microcentrifuge tubes.

### Microbial abundance

To estimate microbial abundance across tissue types, we used dilution plating to estimate the density of colony forming units (CFUs) for each sample. Selective media for growth of fungi (yeast malt agar + chloramphenicol) and bacteria (R2A + cycloheximide) was used. Although not all microbes are culturable, previous work suggests that most dominant species observed to be associated with flowers are culturable on these media types (Morris *et al*., 2020). Plates were incubated for 5 days at 25 °C and colonies counted.

### DNA extraction and sequencing

Genomic DNA was extracted from samples using a ZymoBIOMICS^®^ DNA Microprep kit (Zymo Research, Irvine, CA, USA) at the University of California, Davis (Davis, CA, USA), following the manufacturer’s protocol. Extracted DNA was then sent to the Centre for Comparative Genomics and Evolutionary Bioinformatics at Dalhousie University (Halifax, Nova Scotia, Canada) for library preparation and 16S/ITS amplicon sequencing. Amplicon sequence variants (ASVs) were assigned using DADA2 (Callahan *et al*., 2016). See SI Materials and Methods for information concerning amplification, sequencing, and bioinformatic processing of data.

### Microbial community assembly

To assess effects of nectar composition, two different fungicides, and arrival order on microbiome assembly, we conducted a sequential inoculation experiment. We used three different synthetic nectar environments containing either copper, typically used in organic orchards, or propiconazole, a synthetic, demethylation inhibitor (DMI) often used in conventional orchards. We used microbes commonly observed in nectar, including *Asaia astilbis* (bacterium)*, Aureobasidium pullulans* (yeast)*, Metschnikowia reukaufii* (yeast), and *Neokomagataea thailandica* (bacterium). These species are frequently found on floral tissues and nectar, including almond (Fridman *et al*., 2012; Aizenberg-Gershtein *et al*., 2013; Schaeffer *et al*., 2017). Colonies formed by these species are distinguishable on media. Strains used were isolated from almond nectar or *Epilobium canum* (Onagraceae), a perennial herb native to the foothills of California (Morris *et al*., 2020). Yeast and bacteria strains were cultured on YMA and R2A, respectively, and grown at 25 °C. After three days of growth, microbial cell suspensions for inoculation were diluted to ca. 400 cells μL^−1^ using a hemocytometer just prior to the beginning of the experiment described below.

To prepare the synthetic nectar environments, fungicides were added to filter-sterilized 15% (w/v) glucose:fructose solution supplemented with 0.32 mM amino acids from digested casein (Vannette & Fukami 2014). Fungicide concentration (7500 ppb) was parameterized based on residue analyses of almond flowers and those of other flowering crops (Frazier *et al*., 2015). In each nectar environment, we assessed the strength of priority effects following a two-way, full-factorial design, with three different orders of species introductions (e.g., Tucker and Fukami 2014). Our introduction treatment groups included: (i) simultaneous introductions of the two yeast and bacterial species to the artificial nectar on day 0; (ii) ‘yeast-first’ sequential introductions, in which the two yeast species were introduced first (day 0) and then, 48 h later, the two bacterial species; and (iii) ‘bacteria-first’ sequential introductions, in which the two bacterial species were introduced first (day 0) and then, 48 h later, the two yeast species. The experiment was performed in 200 μL polymerase chain reaction (PCR) tubes (ThermoFisher Scientific Corp.) and lasted for 4 d, which approximates the lifespan of an almond flower (Yi *et al*., 2006). To each tube, we added 9 μL of synthetic nectar at the start, and 0.5 μL suspensions of each respective microbial species (ca. 200 cells) for the appropriate day and treatment combination. We chose this timing as 48 h as a realistic time interval between microbial immigration events in floral systems with displays open for such a duration (Peay *et al*., 2012; Vannette and Fukami 2014). Four days after the first inoculation the experiment was ended and nectar from each tube was divided for chemical analysis and to determine microbial abundance. All treatment combinations were performed in each of the three nectar environments (*N* = 16 per treatment combination).

### Pollination

#### Pollination Experiment One

To test consequences, including non-additive effects, of flower exposure to fungicides and microbes on pollinator foraging behavior, we performed a field assay. Briefly, artificial flowers (Figure S2) designed to mimic those of almond were set out in an array near an apiary at the Harry H. Laidlaw Jr. Honey Bee Research Facility (Davis, CA). Flowers were treated with 200 μL of artificial nectar (same as above) in a fully-crossed design, with three levels for each treatment. For fungicides, treatment levels were: (1) no fungicide (control), (2) organic (copper, 7500 ppb), or (3) conventional (propiconazole, 7500 ppb). With respect to nectar-inhabiting microbes, treatments were: (1) no bacterium or yeast (control), (2) *N. thailandica* (bacterium), or (3)*M. reukaufii* (yeast). Experimental arrays were set ~1–2 m from the hives at the apiary in the morning each day the experiment was performed. Two hours after the start of the experiment each day, the remaining nectar from each flower’s tube was capped, brought back to the laboratory, and weighed to estimate changes in volume. This assay was performed four times (*N* = 12 flowers per treatment combination per assay). See SI Materials and Methods for full experimental details.

#### Pollination Experiment Two

We performed an *in vivo* field assay at an orchard (Zamora, CA, USA) to test for consequences, including non-additive effects, of flower exposure to fungicides and microbes on the quality of pollination services. Briefly, fungicide/microbe treatments mirrored those used in the first pollination assay, with treatment identity randomized among 9 unvisited flowers within an individual tree (*N* = 20 ‘Nonpareil’ variety trees, spaced across alternating rows, with five haphazardly selected in each row). Two microliters of treated nectar was applied to each flower, and after two days of exposure to pollinators, flowers were carefully removed along with the pedicel and placed in individual 1.5 mL microcentrifuge tubes containing 0.5 mL of water. Flowers were positioned such that the stigma did not touch the tube’s surface and the pedicle was in water (Brittain *et al*., 2013). Once returned to the lab, flowers were stored in the dark at room temperature for 72 h to allow pollen tube growth. After this period, pistils were fixed (Farmer’s fixative) and then stored at 4 °C until further processing. Pollen tube growth was assessed using a staining and microscopy procedure, following a previously established protocol (Brittain *et al*., 2013). See SI Materials and Methods for full experimental details.

### Statistical analyses

All analyses were performed in R v.4.0.2 (R Core Team, 2013). We fit linear mixed-effect models with the *lme4* package (Bates *et al*., 2014) to assess the impact of orchard management, and other measured variables on microbial abundance (log-10 transformed CFU counts) and diversity (ASV richness and Shannon diversity index). For each model and response variable examined, management, bloom stage (early/peak), site (edge/interior) and a three-way interaction among them, along with amount of semi-natural habitat surrounding orchards and distance to the nearest apiary were included as predictors, with orchard identity as a random effect to account for repeated sampling. Bacterial and fungal data were analyzed separately for each floral tissue examined. Once fit, we used backward stepwise model selection in the *lmerTest* package (Kuznetsova *et al*., 2017) to identify the best-fit model for each response variable examined. Fixed model terms were retained based on log-likelihood ratio tests, with significance of each calculated using *F* tests, based on Satterthwaite approximation for denominator degrees of freedom (Kuznetsova *et al*., 2016). Finally, to determine if the relative abundance of individual ASVs responded to orchard variables of interest, we used *DESeq2* with Benjamini-Hochberg corrections for multiple testing (Love *et al*. 2014). Orchard management, bloom stage, and apiary distance were treated as predictors in separate models.

Pairwise dissimilarities between fungal and bacteria communities were calculated using the Bray–Curtis dissimilarity metric. We also calculated abundance-weighted UniFrac distances for bacteria; the UniFrac metric uses phylogenetic information to calculate dissimilarities between communities, and was weighted by the relative abundance of ASVs within a sample (Lozupone and Knight, 2005). Finally, we used permutational multivariate analysis of variance (PERMANOVA) to assess the contribution of management, bloom stage (early/peak), site (edge/interior) and all (2- and 3-way) interactions among them, along with amount of semi-natural habitat surrounding orchards and distance to the nearest apiary on community composition. This analysis was performed using *vegan*, based on 1000 permutations (Oksanen *et al*., 2015).

To assess effects of fungicide identity and immigration order on microbiome assembly in our *in-vitro* experiment, we fit linear mixed-effect models for each species with abundance (CFUs) as the response variable and immigration order and fungicide treatment as fixed factors, including their interaction.

For the behavior assay (*Pollination Experiment One*), we fit a linear mixed-effects model with nectar remaining as the response, fungicide and microbe treatments as fixed factors, as well as their interaction. Trial number was included as a random effect. For the in-orchard pollination service assay (*Pollination Experiment Two*), we also fit linear mixed-effects models with pollen germination and number of tubes as response variables, fungicide and microbe treatments as fixed factors, as well as their interaction. Tree identity was included as a random effect.

## Results

### Microbial abundance and diversity

Floral microbial abundance and diversity increased from early to peak bloom across all tissue types, and management strategies, for both bacteria and fungi (Tables S1-S3). The magnitude in which microbial abundance increased, however, varied considerably among these factors. Culturable bacterial CFU abundance from petals (*F_1,53_* = 645.35, *P* < 0.0001), anthers (*F_1,43.92_* = 466.26, *P* < 0.0001), and nectaries (*F_1,54_* = 929.91, *P* < 0.0001) was eleven-, five-, and eight-fold higher at peak bloom, respectively (Figure 2; Table S1) than at the initial sampling.

Like bacteria, fungal CFU abundance (Table S1) also increased from early to peak bloom, increasing two-fold for petals (*F_1,46.03_* = 6.38, *P* = 0.02), two-fold for anthers (*F_1,52_* = 70.97, *P* < 0.0001), and three-fold for nectaries (*F_1,44.04_* = 41.11, *P* < 0.0001).

**Figure 2.**
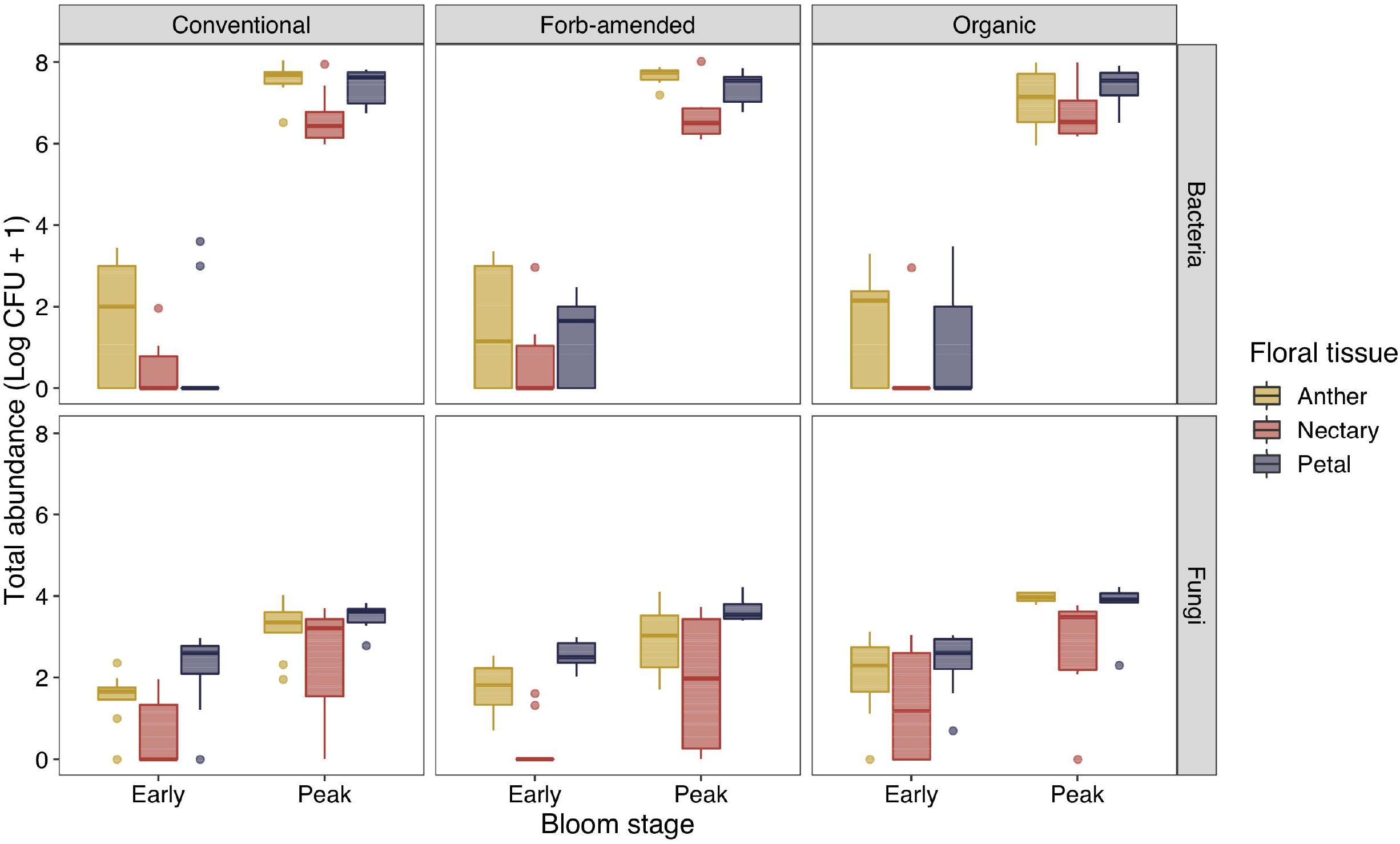
Abundance of a) colony-forming units (CFUs) on R2A media (bacteria) and b) CFUs on YMA (fungi) associated with floral tissues (Anther, Nectary, or Petal) of almond. Flowers were sampled at two stages of bloom (Early or Peak) from orchards that employed different management schemes (Conventional, Forb-amended, or Organic).

Beyond bloom progression, additional orchard factors affected microbial abundance, but in disparate ways for bacteria and fungi across tissue types. Fungi associated with anthers were 27% more abundant in organic orchards than those that were forb-amended or conventional in management practice (*F_2,52_* = 4.61, *P* = 0.01). Moreover, regardless of management scheme, fungi associated with petals were 11% more abundant along the edges of orchards compared to the interior (*F_1,46.03_* = 6.38, *P* = 0.02). However, no effect of sampling location within orchards was detected for bacteria, nor for the amount of natural habitat in the surrounding landscape on the abundance of either bacteria or fungi.

Bacterial diversity increased as bloom progressed for communities associated with petals (richness: *F_1,54_* = 8.85, *P* < 0.01; Shannon index: *F_1,54_* = 6.05, *P* = 0.02), anthers (richness: *F_1,35.60_*= 25.99, *P* < 0.0001; Shannon index: *F_1,51_* = 33.28, *P* < 0.0001), and nectaries (richness: *F_1,40.32_* = 102.96, *P* < 0.0001; Shannon index: *F_1,40.17_* = 25.45, *P* < 0.001) (Figure 3; Tables S2 and S3). All tissues were dominated by Proteobacteria (Figure 4), particularly members of the Pseudomonadaceae, with two taxa being enriched from early to peak bloom overall (BactSeq2: *Pseudomonas* sp., log2-fold change = 2.50, *P_adj_* < 0.001; BactSeq10: *Pseudomonas* sp., log2-fold change = 3.64, *P_adj_* < 0.01). Tree proximity to apiaries within an orchard was found to be associated with bacterial diversity, although the association was weak (Figure S2A): Shannon diversity was higher on flowers closer to apiaries than those that were collected further away, though a significant effect was only detected for anthers (*F_1,51_*= 7.39, *P* < 0.0001) in our models.

**Figure 3.**
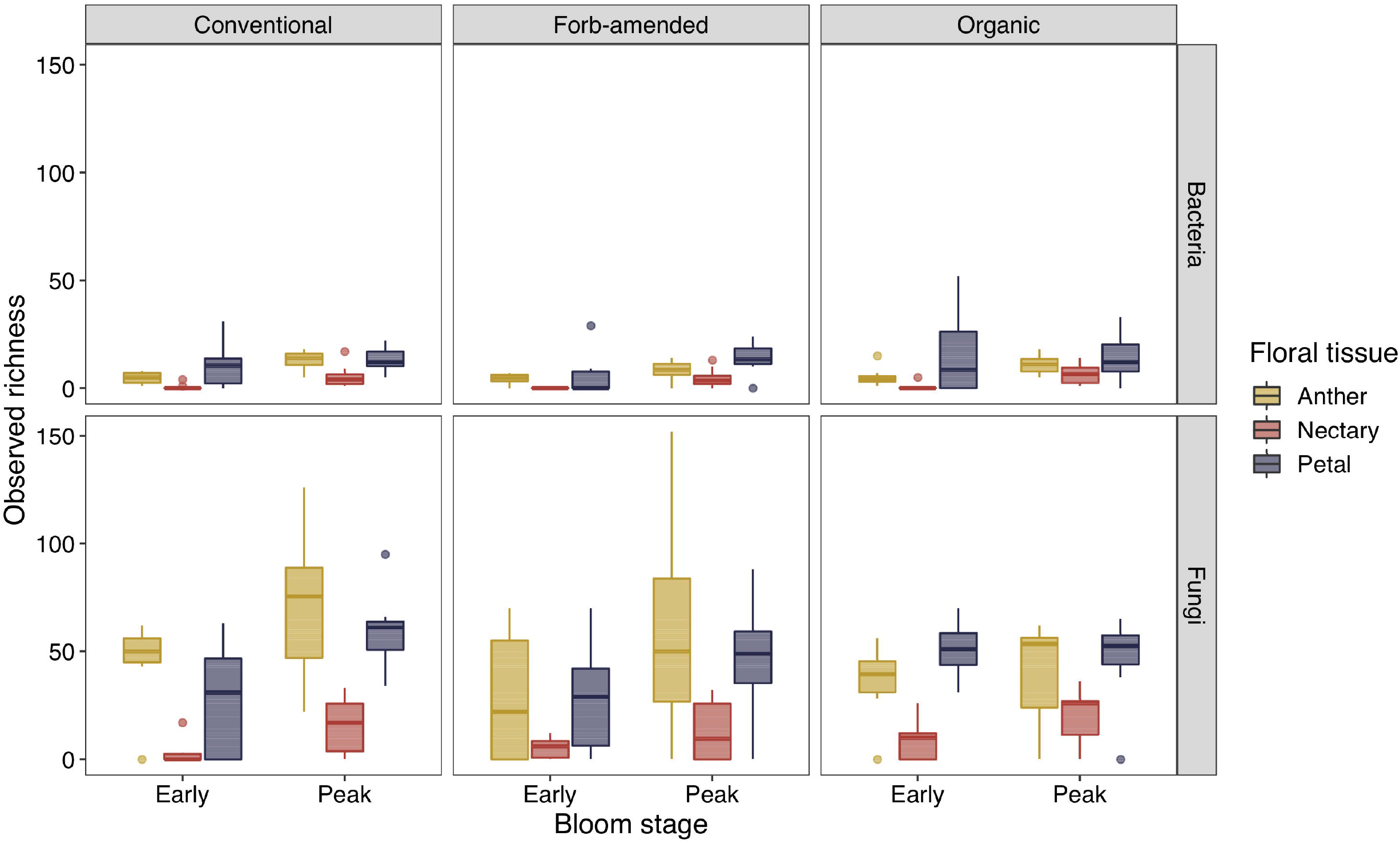
Observed sequence variant richness of a) bacteria and b) fungi associated with floral tissues (Anther, Nectary, or Petal) of almond flowers. Flowers were sampled at two stages of bloom (Early or Peak) from orchards that employed different management schemes (Conventional, Forb-amended, or Organic).

**Figure 4.**
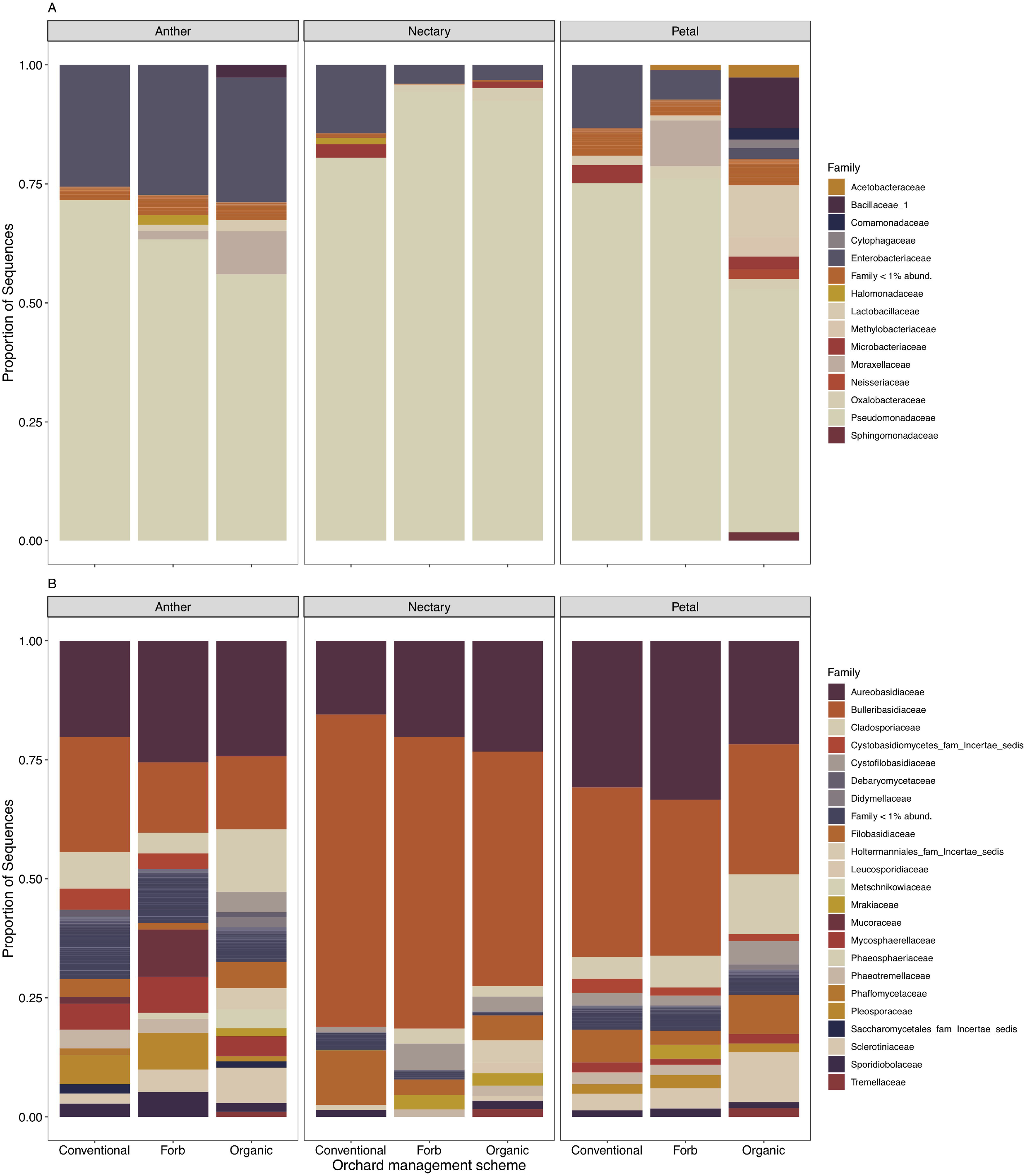
Average relative abundance (Proportion of Sequences) of a) bacterial and b) fungal families associated with floral tissues (Anther, Nectary, or Petal) of almond. Flowers were collected from orchards that vary in management scheme (Conventional, Forb-amended, or Organic).

With the exception of organic orchards, fungal ASV richness generally increased over bloom (Figure 3). This trend was significant for nectaries (richness: *F_1,42.37_* = 6.81, *P* = 0.01) and petals (richness: *F_1,41_* = 5.32, *P* = 0.03). Shannon diversity followed a similar pattern, with a significant increase observed for nectaries (*F_1,42.44_* = 10.22, *P* < 0.01) and petals; however, for the latter this effect depended on orchard management scheme (Management × Bloom stage: *F_2,39_* = 3.50, *P* = 0.04). Specifically, fungal diversity (Shannon index) increased in both conventional and forb-amended orchards by 59% and 12% respectively, while decreasing in organic orchards by 26%. Other orchard-level variables examined, including apiary distance, natural habitat in the surrounding landscape, and site sampled within orchards, had no effect on observed fungal ASV richness or Shannon diversity (Tables S2 and S3). We also detected no correlation between apiary distance and either diversity metric (Figure S2b); however, DESeq2 analyses revealed *Vishniacozyma carnescens* (syn. *Cryptococcus carnescens*) ASVs that significantly declined in abundance with tree distance from apiaries (FunSeq13: log2-fold change = −0.02, *P_adj_* < 0.001; FunSeq58: log2-fold change = −0.02, *P_adj_* < 0.01).

Fungal communities associated with floral tissues were generally dominated by members of the Aureobasidiaceae and Bulleribasidiaceae (Figure 4), including *Aureobasidium pullulans, V. victoriae* (syn. *C. victoriae*), and *V. carnescens*. Over bloom, *A. pullulans* in particular was found to significantly increase in relative abundance (FunSeq1: log2-fold change = 2.96, *P_adj_* < 0.0001), along with *Gelidatrema spencermartinsiae* (syn. *C. spencermartinsiae*, FunSeq16: log2-fold change = 2.27, *P_adj_* < 0.01), *Filobasidium wieringae* (syn. *C. wieringae*, FunSeq34: log2-fold change = 3.87, *P_adj_* < 0.0001), and *Buckleyzyma aurantiaca* (syn. *Rhodotorula aurantiaca*, FunSeq26: log2-fold change = 1.14, *P_adj_* < 0.01). Taxa observed to significantly decline in relative abundance included *Cladosporium delicatulum* (FunSeq3: log2-fold change = −1.52, *P_adj_* < 0.001), a widely distributed saprobe species, and *Naganishia friedmannii* (syn. *C. friedmannii;* FunSeq27: log2-fold change = −3.28, *P_adj_* = 0.02).

Bacterial and fungal species composition differed between sampling times (Table 1), with bloom stage explaining 3-21% of variation in composition, depending on flower tissue. Orchard-level predictors (e.g., natural habitat) generally explained less variation in composition (Table 1). Orchard management was not associated with bacterial species composition, but did predict variation in fungal composition in anthers (*R^2^* = 0.073, *F_2,33_* = 2.11, *P* = 0.001) and petals (*R^2^* = 0.078, *F_2,33_* = 2.60, *P* = 0.005). The amount of semi-natural habitat in the surrounding landscape, as well as apiary distance, were both generally found to be associated with shifts in bacterial and fungal community composition, with a particularly notable effect of apiary distance on bacteria in nectar (*R^2^* = 0.176, *F_2,20_* = 5.65, *P* = 0.001).

**Table 1.**
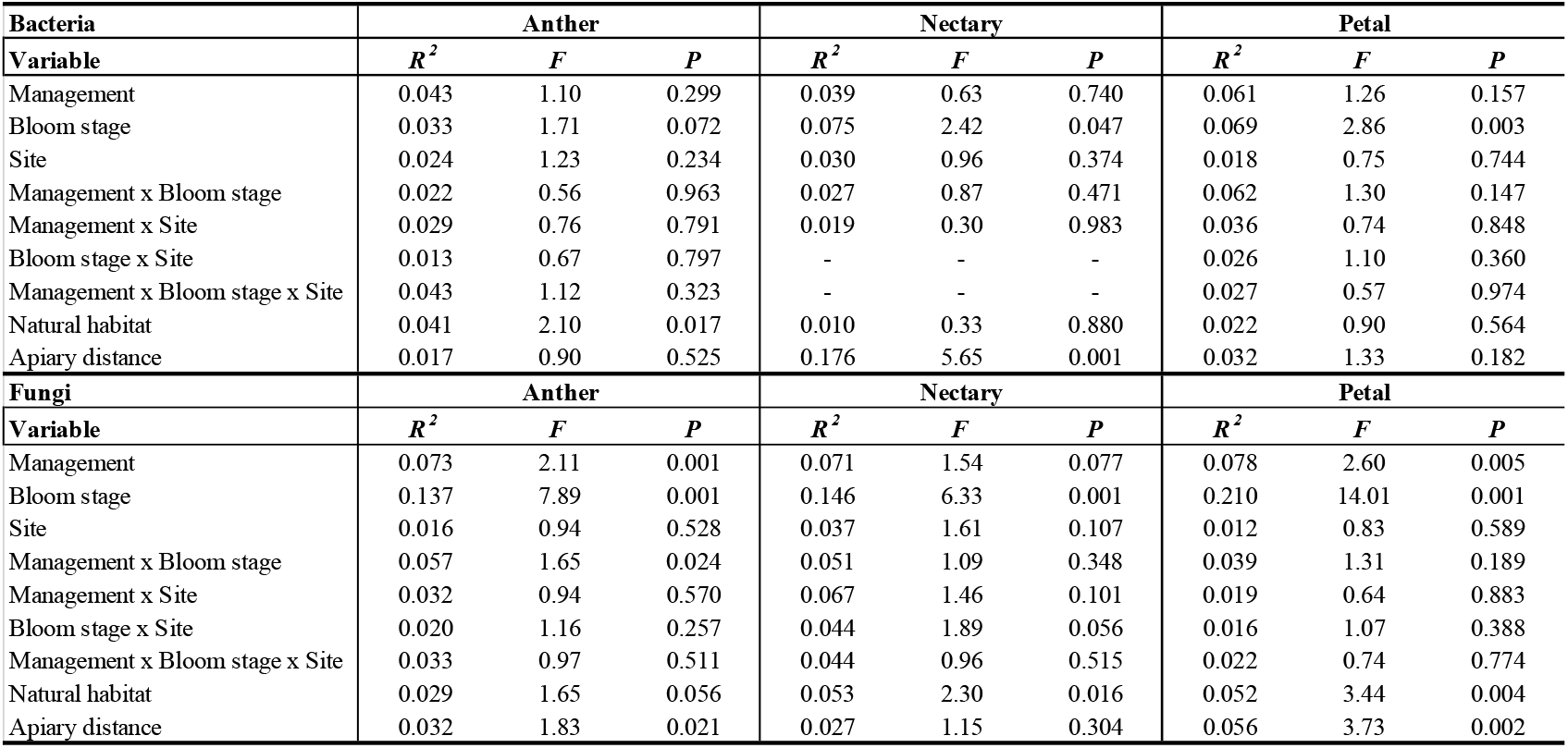
PERMANOVA results of Bray-Curtis dissimilarity between bacterial and fungal communities associated with almond floral tissues.

### Microbial community assembly in floral nectar

In the in-vitro sequential inoculation experiment, arrival order significantly affected species’ densities in nectar. However, this effect depended on the nectar environment and presence of fungicide residues (Figure 5). The bacterium *Asaia* was the only species that could persist across all nectar environments (*F_2,134_* = 2.64, *P* = 0.08). Interestingly, the bacterium *Asaia* reached higher densities when introduced after or simultaneously with yeast species (*F_2,134_* = 22.89, *P* < 0.0001). Of the yeasts, persistence and growth were strongly dependent on the nectar environment: *Metschnikowia* grew best in control nectar (*F_2,134_* = 31.63, *P* < 0.0001), while *Aureobasidium* performed better in the copper treatment (*F_2,134_* = 16.94, *P* < 0.0001). Finally, propiconazole had strong inhibitory effects on the growth of both species.

**Figure 5.**
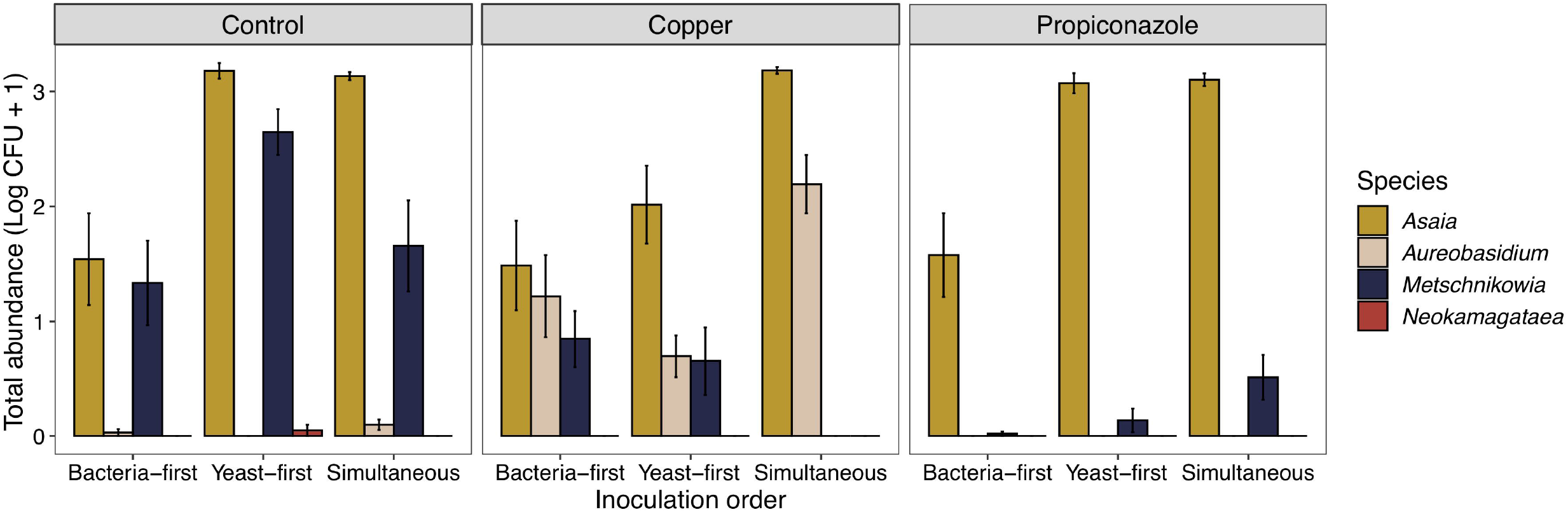
Mean abundance of colony-forming units (CFUs) of bacteria (gold = *Asaia*, red = *Neokamagataea*) and yeasts (tan = *Aureobasidium*, blue = *Metschnikowia*) when introduced in different orders (Bacteria-first, Yeasts-first, or Simultaneously) to nectar environments treated with fungicides (Copper or Propiconazole at 7500 ppb). Simultaneous introductions were carried out on day 0, while sequential introductions on days 0 and 2.

**Figure 6.**
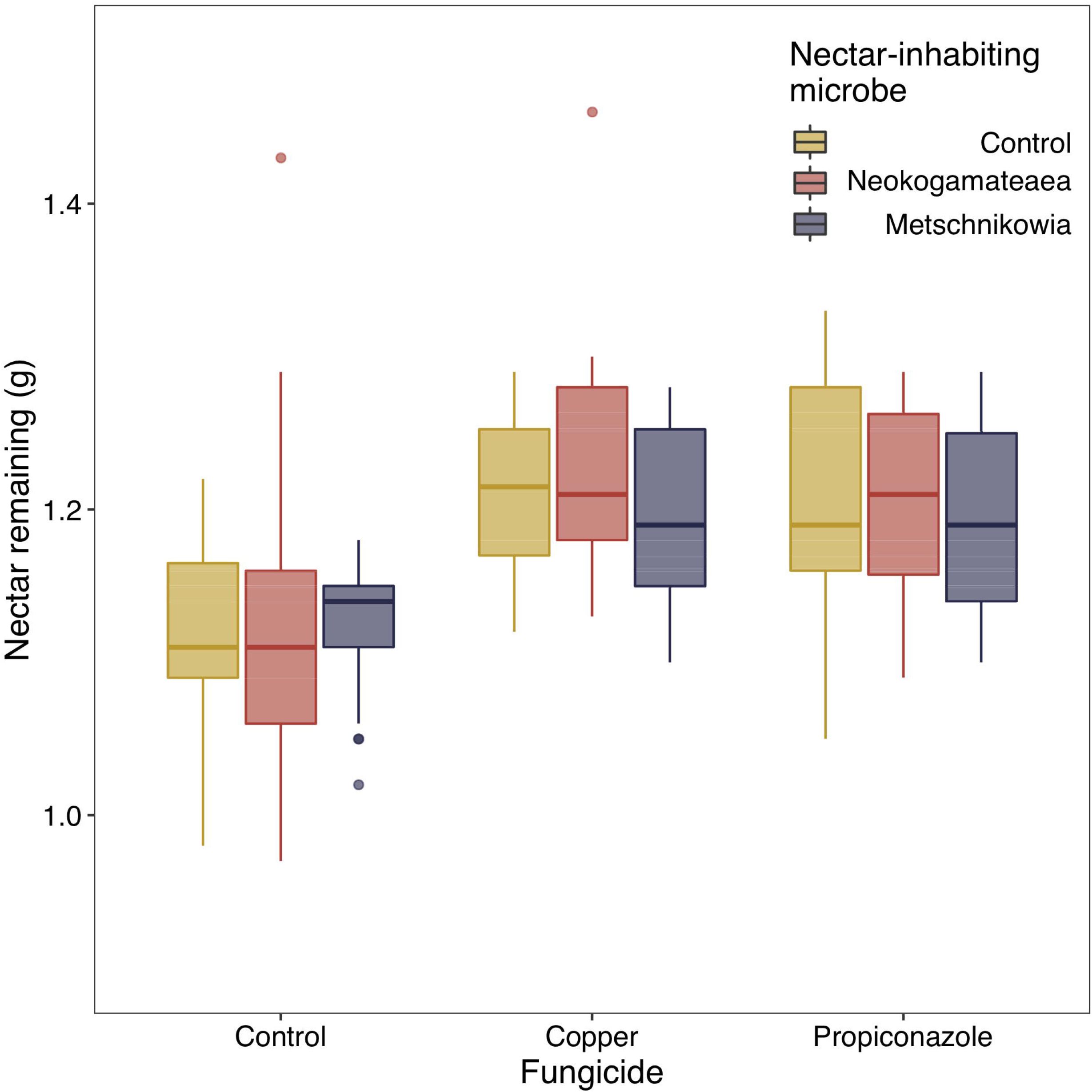
Nectar remaining (g) in artificial almond flowers treated with fungicides (Copper or Propiconazole) and nectar-inhabiting microbes (red = bacterium *Neokogamataea thailandica*, blue = yeast *Metschnikowia reukaufii*).

### Microbial and fungicide effects on honey bee foraging and pollination

Fungicides reduced the amount of nectar removed by honey bee foragers (*F_2,311_* = 130.76, *P* < 0.0001). In contrast, neither microbe affected forager nectar removal (*F_2,311_* = 1.90, *P* = 0.15), nor did we detect a significant interaction between microbial inoculation and fungicide treatment (*F_4,311_* = 2.19, *P* = 0.07). In the orchard experiment, neither fungicide application nor nectar-inhabiting microbes affected pollen germination or pollen tube number (Table S5).

## Discussion

The results presented here demonstrate that orchard management practices can mediate flower microbiome structure, though the magnitude of these effects can hinge on a variety of factors. Of those examined, timing of bloom was the most consistent predictor: bacterial and fungal abundance and diversity were higher at peak bloom than at the start across all floral tissue types and management strategies. Sampling time with respect to bloom intensity has been documented previously (e.g., Smessaert *et al*., 2019), and may be driven by variation in a combination of interrelated variables, including temperature, pollinator activity, and host metabolism. However, other orchard-level variables affected microbial abundance and diversity, with host proximity to apiaries and orchard management having significant effects on both bacteria and fungi, respectively.

The results described here support a key role for pollinator foraging in the assembly of the almond flower microbiome. For example, the orchard survey detected a strong signature of tree proximity to apiaries on epiphytic bacterial diversity of almond flower nectaries and reproductive structures, as both bacterial richness and diversity decreased the further trees were from apiaries. Within orchards, honey bees forage more intensively near their apiary (Bates *et al*., 2014), dispersing bacteria among flowers (Vannette and Fukami, 2018; Russell *et al*., 2019), and generate spatial variation in microbial transmission and resulting flower microbial communities. Moreover, while foraging on flowers, individual honey bees can display consistent behaviors, including focus on either pollen and nectar collection only, affecting the degree of contact with intrafloral tissues (Bosch and Blas, 1994; Thomson and Goodell, 2001). Finally, although contact with petals can occur as foragers side-work for nectar (Thomson and Goodell, 2001), these collective behaviors may explain the lack of a significant effect of apiary distance on bacterial diversity observed on petals vs. nectar and reproductive structures of the flower. In sum, these patterns point to increased consideration of both apiary spacing, and pollinator foraging behavior, as interest grows in leveraging pollinators as vectors of microbial biocontrol agents to combat disease (Kevan *et al*., 2008; Menzler-Hokkanen and Hokkanen, 2017).

Pollinator foraging could also affect microbial arrival to flowers and community assembly, generating priority effects between microbial species (Peay *et al*., 2011; Vannette and Fukami, 2014). Indeed, in our laboratory experiment, immigration history had a significant effect on species’ persistence and densities in nectar. These effects, however, were dependent on both the biotic environment and presence of fungicides. At the concentrations tested, both fungi were highly susceptible to the synthetic fungicide propiconazole, while *Aureobasidium* was much less so in comparison to *Metschnikowia* when challenged with copper. Surprisingly, the bacterium *Asaia* persisted in all nectar environments, while *Neokomagataea* did not, in contrast to previous findings (Tucker and Fukami, 2014). Additional work on the sensitivity of these taxa to fungicides across different sugar environments may explain these patterns. Taken together, these results suggest that arrival order and environmental filtering via the presence of fungicides can potentially affect beta diversity among nectar microbial communities in managed flowering systems. As nectar-inhabiting fungi and bacteria can differ in their effect on nectar chemistry (Vannette and Fukami, 2018), it is possible that microbial effects stemming from differing community compositions may manifest to affect the quality of pollination services provided among managed flowering systems.

Honey bee foraging was affected by fungicides, but not by nectar-inhabiting microbes, in contrast to previous work (Vannette *et al*., 2013; Good *et al*., 2014; Rering *et al*., 2018). Regardless of fungicide type, honey bees removed less nectar from artificial flowers with copper and propiconazole residues, demonstrating that fungicide application and residual contamination of floral rewards can affect forager decisions. Exposure to fungicide-contaminated rewards can affect pollinator health and pollination in agroecosystems, including almond. Such consequences range from negative effects on larval development, pollinator cognition, and even mortality (Johnson, 2015), as has been observed with honey bee workers directly exposed to a range of fungicides commonly employed for pathogen control in almonds. Although fungicide residues in nectar deterred honey bees, these shifts in foraging behavior did not translate to noticeable effects on pollination services in almonds measured through pollen germination and tube number. Honey bees foraging on almond flowers however often alternate between foraging for nectar or pollen (Bosch and Blas, 1994). These nectar and pollen foragers differ markedly in the quality of services that they provide, with pollen foragers being on average five times more effective in affecting fruit set than those foraging for nectar (Bosch and Blas, 1994). Given that our flowers were exposed for a long enough duration to allow pollinators to forage for both resources, those that foraged for pollen alone likely conferred adequate pollination observed in our experiment.

Our results highlight multiple orchard management practices that can shape the assembly of crop-associated microbiota during flowering and pollination. We documented temporal changes in microbial abundance and composition, but also detected effects of managed pollinators and natural areas, suggesting a key role of immigration in determining species composition in many floral tissues. Combined with the potential for agrochemicals to differentially affect microbial growth and species interactions, we outline a few factors that likely contribute to flower microbiome assembly. Because flowers form the template for potential reproductive output that is translated through interactions with pollinators, understanding linkages between management, the assembly of the floral microbiome and its impact on pollination in crops can reveal how microbial interactions affect both crop yield and quality.

## Authors’ contributions

RS conceived the idea for the study and collaborated with DC, JB, TF, NW, and RV in designing the survey and experiments performed. RS carried out the survey, experiments, analyses, and wrote the first draft of the manuscript. DC and JI contributed ArcGIS data and analysis. All authors contributed to revisions.

## Acknowledgements

We are grateful to the growers for permission to work on their properties, and E. Niño for access to an apiary. We also thank E. Bloom, S. Cibotti, L. Hack, G. Hall, W. Menendez, and H. Pathak for field and laboratory assistance. This work was supported by a USDA ELI Postdoctoral Fellowship (2017-67012-26104) awarded to RS, USDA NIFA (grant accession nos 1003539 and 1014754) to DC, Project Number 6036-2200-028-00-D for JB, and UC Davis start-up and Hatch NE1501 funds awarded to RV.

