## Supplemental Information for "Ecological dynamics of the almond floral microbiome in relation to crop management and pollination"

**Supporting Information – Material and Methods**

***DNA extraction, sequencing, and bioinformatic processing***

Genomic DNA was extracted from samples using a ZymoBIOMICS® DNA Microprep kit (Zymo Research, Irvine, CA, USA) at the University of California, Davis (Davis, CA, USA), following the manufacturer’s protocol. Extracted DNA was then sent to the Centre for Comparative Genomics and Evolutionary Bioinformatics at Dalhousie University (Halifax, Nova Scotia, Canada) for library preparation and 16S/ITS amplicon sequencing. Amplicon fragments were PCR-amplified from DNA in duplicate using separate template dilutions (1:1 & 1:10) and high-fidelity Phusion polymerase (New England BioLabs Inc., Ipswich, MA, USA). A single round of PCR was performed using "fusion primers" (Illumina adaptors + indices + specific regions) targeting either the 16S V4-V5 (bacteria) or ITS2 (fungi) regions with multiplexing. PCR products were verified by running a high-throughput Invitrogen 96-well E-gel (Thermo Fisher Scientific Corp., Carlsbad, CA, USA). Any samples with failed PCRs (or spurious bands) were re-amplified by optimizing PCR conditions to produce correct bands to complete a sample plate before continuing with sequencing. The PCR reactions from the same samples were pooled in one plate, cleaned, and normalized using high-throughput Invitrogen SequalPrep 96-well Plate Kit (Thermo Fisher Scientific Corp.). Amplicon samples were then run on an Illumina MiSeq using 300+300 bp paired-end V3 chemistry. Raw sequences are available on the NCBI Short Read Archive (SRA) under BioProject PRJNA660037.

Demultiplexed sequences were trimmed of trailing low-quality bases using the *DADA2* pipeline (v.1.6.0; Callahan *et al*., 2016). Paired-end reads were quality-filtered, error-corrected, and assembled into amplicon sequence variants (ASVs). Chimeras were detected, removed, and taxonomic information assigned to each ASV using the RDP Naïve Bayesian Classifier (Wang et al. 2007), trained to either the RDP training set (v.14) or UNITE general fasta release (v.7.2) for bacteria or fungi respectively. 16S rRNA gene sequences were aligned using *DECIPHER* (Wright 2016), and a phylogenetic tree was generated from the filtered alignment using *phangorn* (Schliep 2011). ASVs that failed to classify to kingdom or were identified as chloroplast or mitochondrial sequences were discarded. Moreover, potential contaminant ASVs were identified through inclusion of negative controls during sample and sequence processing, and removed using the ‘prevalence’ method with the *decontam* package (Davis *et al*., 2017). This filtering resulted in samples sequenced at a mean depth of 17203 sequences per sample for bacteria and 29,728 sequences per sample for fungi. Samples were then rarefied to an even depth (bacteria: 287; fungi: 1328), allowing us to retain nearly all of the dataset (excluding bacterial samples: AbeleASEdgeR1, BeemanNEdgeR1, LongASIntR1; fungal samples: GallConASIntR1, GallForbASIntR1). Though these read cutoffs are comparatively low, sampling curves indicate that we were able to identify the majority of taxa present in samples (Figure S1).

***Pollination experiments***

***Pollination Experiment One –*** To test consequences, including non-additive effects, of flower exposure to fungicides and microbes on pollinator foraging behavior, we performed a field assay. Briefly, artificial flowers designed to mimic those of almond were set out in an array near an apiary at the Harry H. Laidlaw Jr. Honey Bee Research Facility (Davis, CA). Flowers were constructed from white cardstock and 1.5 mL centrifuge tubes (Figure S3). The array consisted of 12 green-painted bamboo sticks (~1m tall), with flowers fixed to each stick using florist’s wire and green florist’s tape. Each stick held 9 flowers, with each assigned to a nectar treatment (detailed below). The array was placed ~1–2 m away from the hives at the apiary. Prior to the start of the experiment, flowers were filled with filter-sterilized 15% (w/v) glucose:fructose solution supplemented with 0.32 mM amino acids from digested casein. Nectar was replenished daily for a period of three days to allow honey bee foragers to identify and recruit to the array.

Flowers were treated in a fully-crossed design, with three levels for each treatment. For fungicides, treatment levels were: (1) no fungicide (control), (2) organic (copper, 7500 ppb), or (3) conventional (propiconazole, 7500 ppb). These concentrations mirror those used in the microbial community assembly experiment outlined above. With respect to nectar-inhabiting microbes, treatments were: (1) no bacterium or yeast (control nectar), (2) *Neokogataea thailandica* (bacterium), or (3) *Metschnikowia reukaufii* (yeast). When preparing the microbe-inoculated synthetic nectar, all preparations were incubated at 25°C for 3 days prior to each day of each experiment in filter-sterilized 15% (w/v) glucose:fructose solution supplemented with 0.32 mM amino acids from digested casein. Individual colonies of the appropriate species were then diluted to ~400 cells per µl each experimental day. The control nectar was prepared the same way, except that it was not inoculated with microbes. Instead, 1.5 mL of filter-sterilized 15% (w/v) glucose:fructose solution supplemented with 0.32 mM amino acids from digested casein was added.

New sterile vials (and cardstock flowers) containing 200 µl of fresh microbe-inoculated synthetic nectar were used each day. Three of the 12 stakes were bagged with mesh (mesh size: 1 mm) to deny access by bees to account for evaporative effects on nectar weight. Approximately two hours after the start of the experiment each day, the remaining nectar from each flower’s tube was capped, brought back to the laboratory, and weighed using a microbalance to estimate changes in volume. Each day, the experiment began at ~7∶00 AM. Direct observations of the artificial flowers were conducted during the first 30 minutes and the last 15 minutes of each of the 2-hour experimental periods. The experiment was repeated four times.

***Pollination Experiment Two*** – We performed an ­*in vivo* field assay at an orchard (Zamora, CA, USA) to test for consequences, including non-additive effects, of flower exposure to fungicides and microbes on the quality of pollination services. Briefly, fungicide/microbe treatments mirrored those used in the first pollination assay, with treatment identity randomized among 9 unvisited flowers within an individual tree (*N* = 20 ‘Nonpareil’ variety trees, spaced across alternating rows, with five haphazardly selected in each row). Within individual trees, and on separate branches, flowers at the ‘popcorn’ stage of flower development (Yi *et al*., 2006) were chosen for treatment. These flowers had yet to be visited by pollinators, and hence were suitable for potential assessment of pollination service quality, and how it was affected by both fungicides and nectar-inhabiting microbes. Petals were gently pried apart, and 2 µL of treated artificial nectar was applied with pipettor. After two days of exposure to pollinators, flowers were carefully removed along with the pedicel and placed in individual 1.5 mL microcentrifuge tubes containing 0.5 mL of water. Flowers were positioned such that the stigma did not touch the tube's surface and the pedicle was in water (Brittain *et al.,* 2013). Once returned to the lab, flowers were stored in the dark at room temperature for 72 h to allow pollen tube growth. After this period, pistils were fixed (Farmer’s fixative) and then stored at 4°C until further processing.

To examine pollen tube growth, we used an established protocol (Brittain *et al.*, 2013). Briefly, pistils were removed from fixative and then softened by boiling in 5% Na_2_SO_3_ for 30 min. Following boiling, pistils were soaked in tap water for 20 min, then incubated for 24 h in a decolorized solution of 0.1% aniline blue dye, dissolved in 0.1 N K_3_PO_4_ (Currier 1957). Softened, stained pistils were squashed onto individual microscope slides, then observed with an epifluorescent microscope (Nikon Eclipse 80i with a CFL-FITC filter) to count the number of pollen tubes initiating growth on the stigma that reached the base of the style.

bees and honey bees for pollination services. *Proceedings of the Royal Society B: Biological Sciences*, *280*(1754), 20122767.

Callahan, B. J., McMurdie, P. J., Rosen, M. J., Han, A. W., Johnson, A. J. A., & Holmes, S. P.

(2016). DADA2: high-resolution sample inference from Illumina amplicon data. *Nature methods*, *13*(7), 581-583.

Currier, H. B. (1957). Callose substance in plant cells. *American Journal of Botany*, 478-488.

Davis, N. M., Proctor, D. M., Holmes, S. P., Relman, D. A., & Callahan, B. J. (2018). Simple

statistical identification and removal of contaminant sequences in marker-gene and metagenomics data. *Microbiome*, *6*(1), 226.

Schliep, K. P. (2011). phangorn: phylogenetic analysis in R. *Bioinformatics*, *27*(4), 592-593.

Wang, Q., Garrity, G. M., Tiedje, J. M., & Cole, J. R. (2007). Naive Bayesian classifier for rapid

assignment of rRNA sequences into the new bacterial taxonomy. *Applied and environmental microbiology*, *73*(16), 5261-5267.

Wright, E. S. (2016). Using DECIPHER v2. 0 to analyze big biological sequence data in R. *R*

*Journal*, *8*(1).

Yi, W., Law, S. E., Mccoy, D., & Wetzstein, H. Y. (2006). Stigma development and receptivity

in almond (*Prunus dulcis*). *Annals of Botany*, *97*(1), 57-63.
