## Supplementary figures and images for "Ecological dynamics of the almond floral microbiome in relation to crop management and pollination"

### Supplemental Figure 1

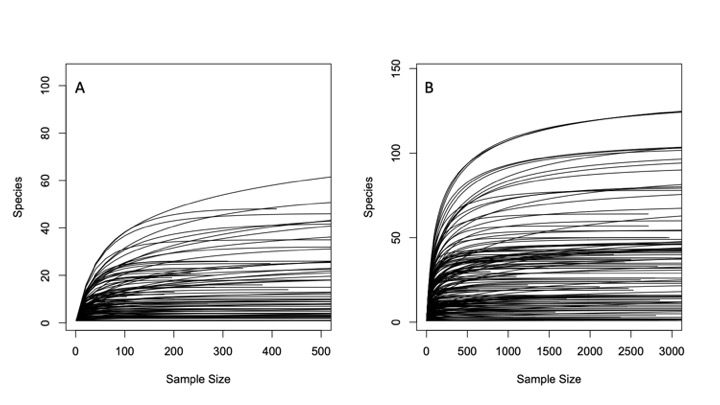

### Supplemental Figure 2

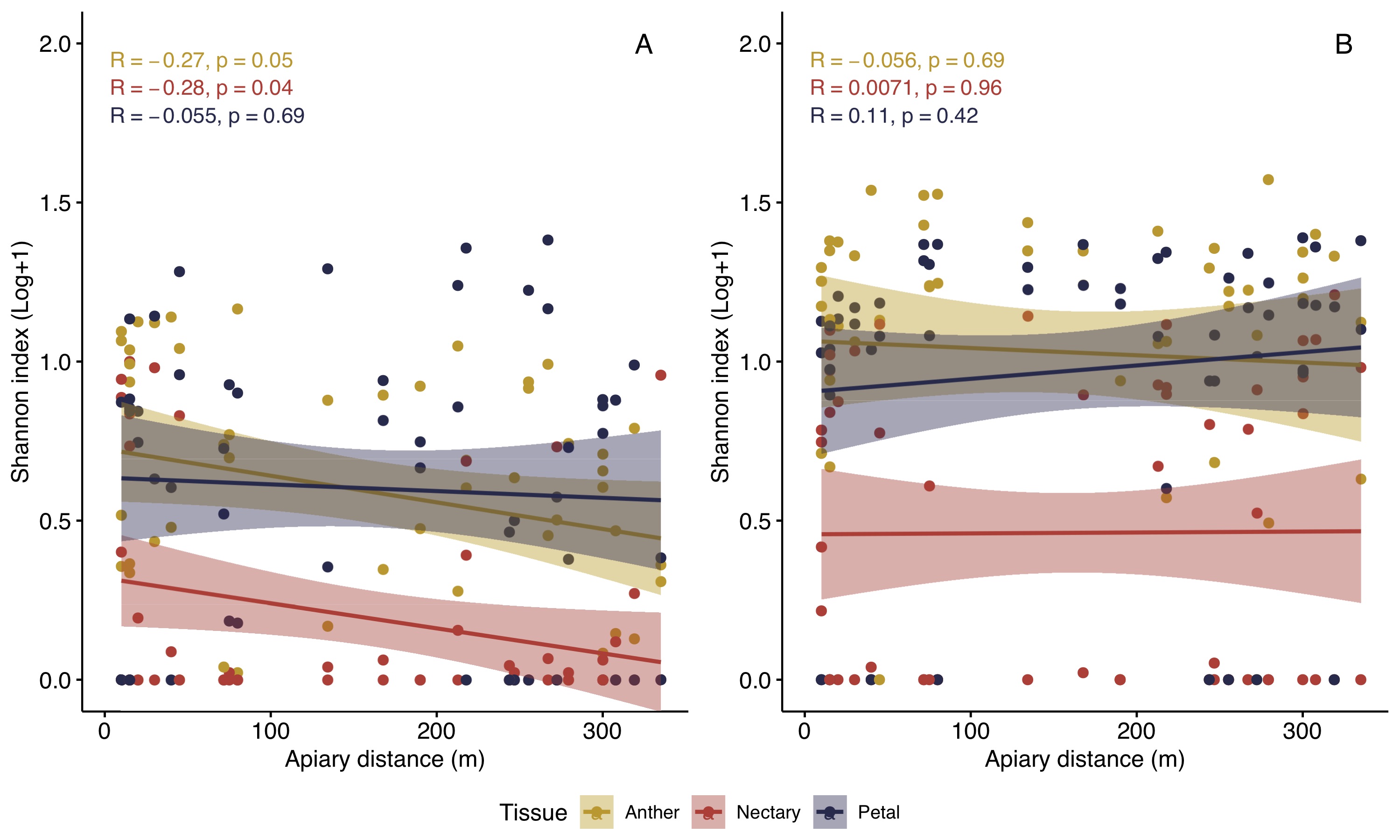

### Supplemental Figure 3

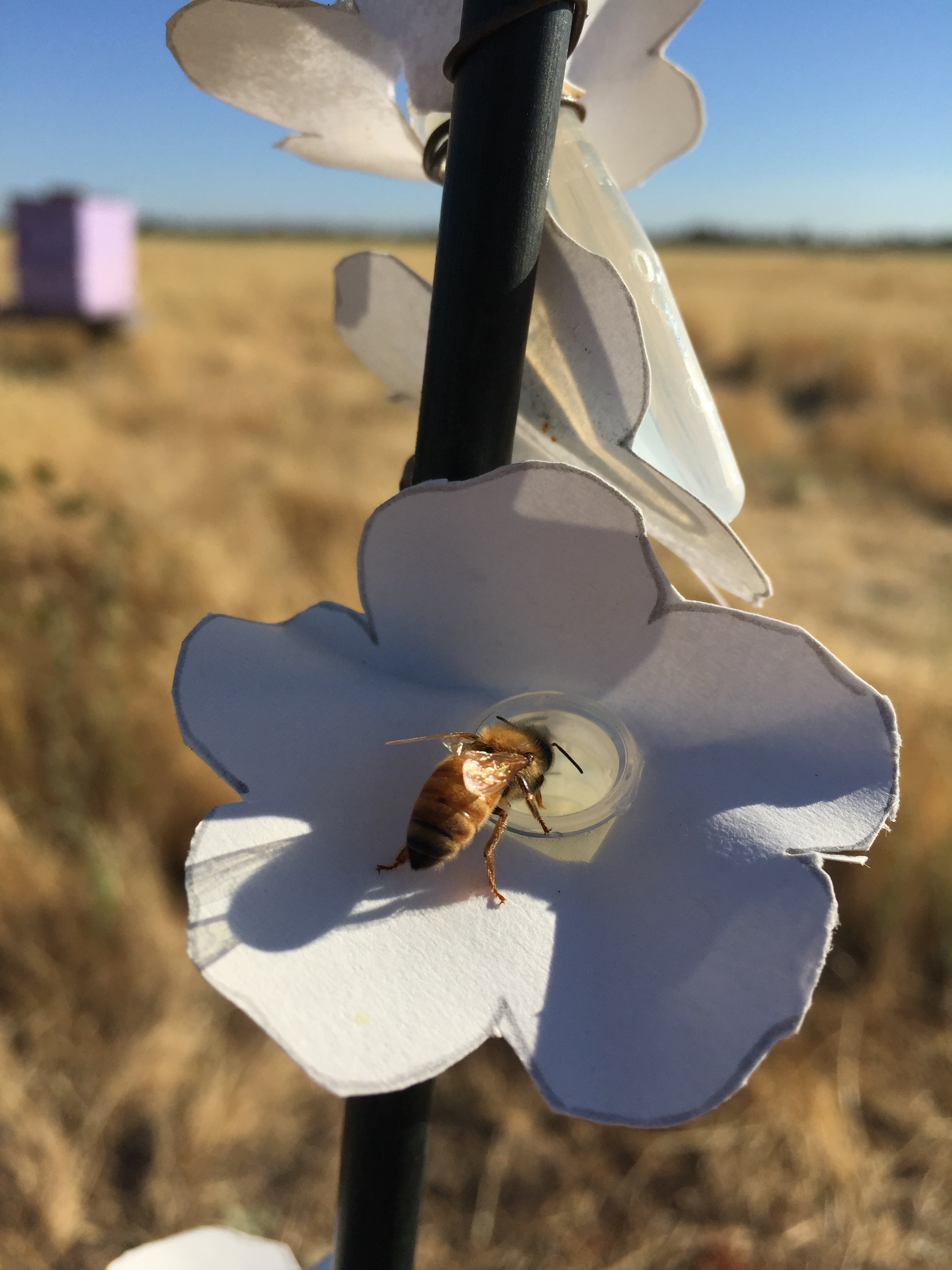
